# The minimal number of genes needed to identify a tumor

**DOI:** 10.1101/2024.07.25.604730

**Authors:** Gabriel Gil, Augusto Gonzalez

## Abstract

We demonstrate that the global state of a Gene Regulatory Network **[1]** may be labeled by a few genes in spite of the fact that there are thousands of genes participating in it. For example, the expression values of only 3 genes are enough to discriminate between a tissue sample coming from a normal lung or a lung adenocarcinoma. We follow a pragmatic procedure, dependent on the sample set, but which is expected to become exact for large enough sets of samples. The proof relies on a scheme for the construction of perfect classification panels of genes **[2]**, inspired by rough set theory **[3]**.

---

The aim at molecularly characterizing a tumor is basically twofold. First, we would like to identify mechanisms implicated in its development **[4]**. And second, there is a natural interest in defining tumor subgroups **[5]**, which could be further used to select the appropriate therapy **[6]**. Both aspects are related to a conceptual question: which is the minimal set of genes needed to identify a tumor? Indeed, if this set is found in a given tissue, for example, any tumor may be labeled by the deregulated genes in the set, and tumor subgroups are naturally defined.

On the other hand, clinical practice states that it is very common to find reduced gene panels whose expression values in body fluids allow a diagnosis with relatively high accuracy **[7]**. These facts point to the conclusion that, in spite of the large amount of genes participating in the Gene Regulatory Network -dozens of thousands- a set of a few genes should be able to discriminate between two of its states: the normal tissue and the tumor.

The situation is illustrated in **Fig. 1** for 12 tissues. Data comes from The Cancer Genome Atlas (TCGA) database **[8]** (https://portal.gdc.cancer.gov/). The data contains expression values for 60,481 genes. In the top panel, we show results from Principal Component Analysis (PCA) for Lung Adenocarcinoma (LUAD) **[9]**. The normal and tumor clouds of samples are well separated along the PC1 direction. We select the main 30 genes along this direction, following a procedure similar to the Page Ranking algorithm **[10]**, and replot the samples along the new PC1 axis. The new median between cloud centers is used to define normal and tumor regions, and thus to roughly reclassify samples, as shown in the center panel. The results are drawn in the bottom panel for LUAD and other 11 tissues, for which the same procedure was applied. In this bottom figure, the *x* axis is the distance between normal and tumor cloud centers for the given tissue in the full space of 60,481 genes **[11,12]**. The *y* axis, on the other hand, is the ratio of false normal samples discovered with the simplified procedure and only the main 30 genes along PC1. As expected, tissues with distant clouds are best classified with only 30 genes. (The exact meaning of used TCGA abbreviations for tumors may be found in **Supplementary Table I**).

**Fig. 1.**
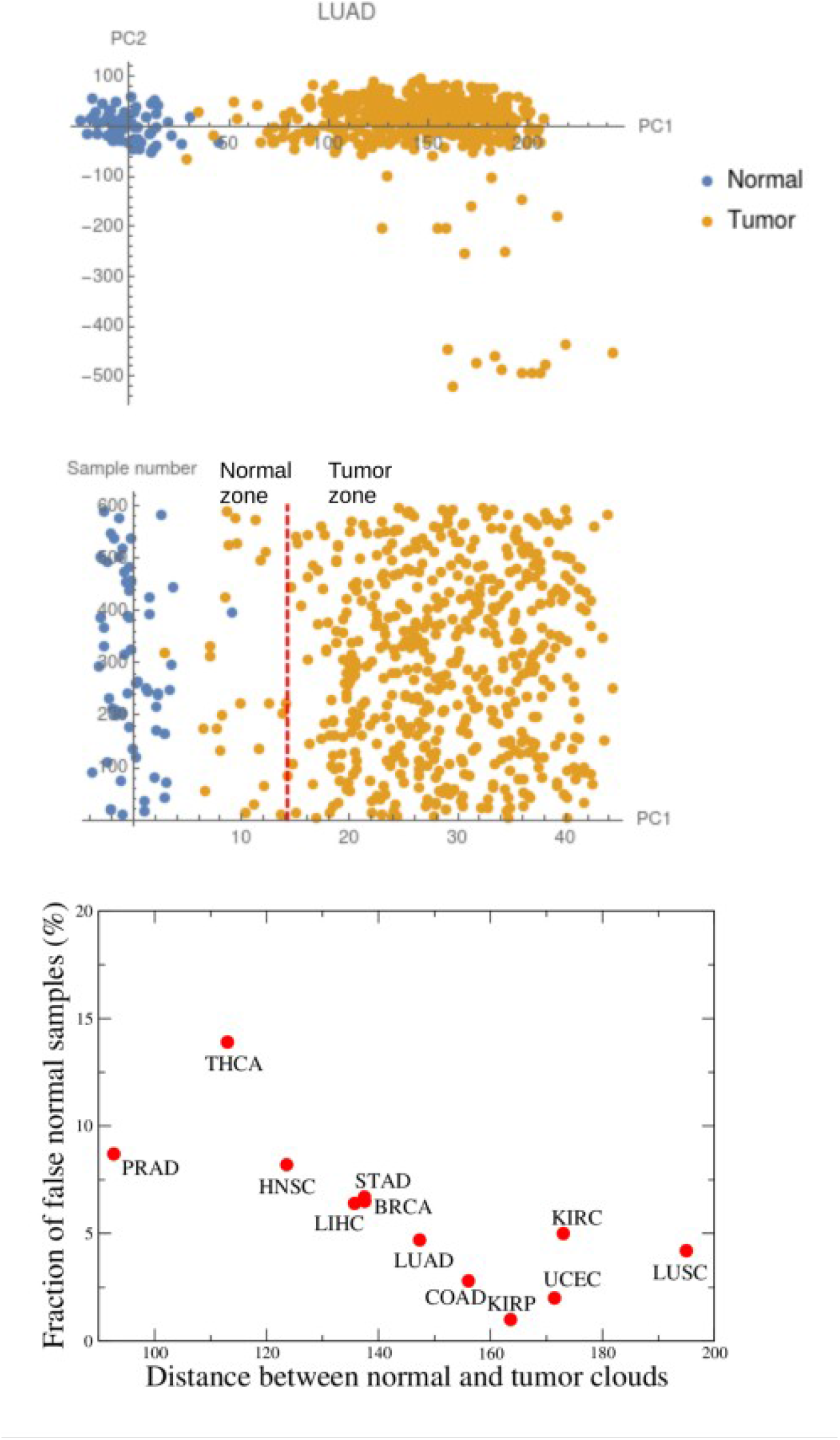
Performance of 30-gene panels in different tumors. **Top panel**: PCA diagram for LUAD. **Center panel**: The distribution of LUAD samples along the PC1 coordinate when only the main 30 genes are considered. The red dashed line is the median between cloud centers, used to reclassify samples. **Bottom panel**: Fraction of false normal samples in different tissues, discovered with the main 30 genes along PC1, as a function of the inter-cloud distance.

The main contribution of the present paper is to show that it is possible to do much better and to get more refined, and even exact, results by following a simple scheme **[2]**. First, define groups or classes of genes according to their expression patterns. As a consequence, inside a group we may define the normal and deregulated expression regions for each gene. Second, conform the binarized table of expression values (0 – normal, 1 – deregulated) for every gene in the sample set. This table is similar to the ones used in applications of rough set theory **[3]**.

We shall illustrate the methodology in a gene set defined to be a “normal concept – only tumors above” class in LUAD. Many other possibilities, and consequently many other gene panels, could be defined. The interested reader may consult **Ref. [2]** for details, explanations and an inventory of gene panels for 12 tissues.

**Fig. 2** shows the distribution of expression values for the PYCR1, ALDH18A1 and TRIM27 genes in lung normal and LUAD samples. We may explicitly define the normal and deregulated expression intervals for each gene. By definition, the expression values for all normal samples are contained in the normal intervals, that is the binarized gene expression values are zero for all of the normal samples. In the tumor subset, however, the binarized expressions could take values 0 or 1, where 1 means that the expression is above the normal interval. Additionally, PYCR1, ALDH18A1 and TRIM27 are deregulated or expressed above the normal threshold in a significant set of tumor samples. Thus, the previous gene set belongs to the “normal concept – only tumor above” class. A deregulated expression value (1) in any of the previous genes is a marker of a tumor.

**Fig. 2.**
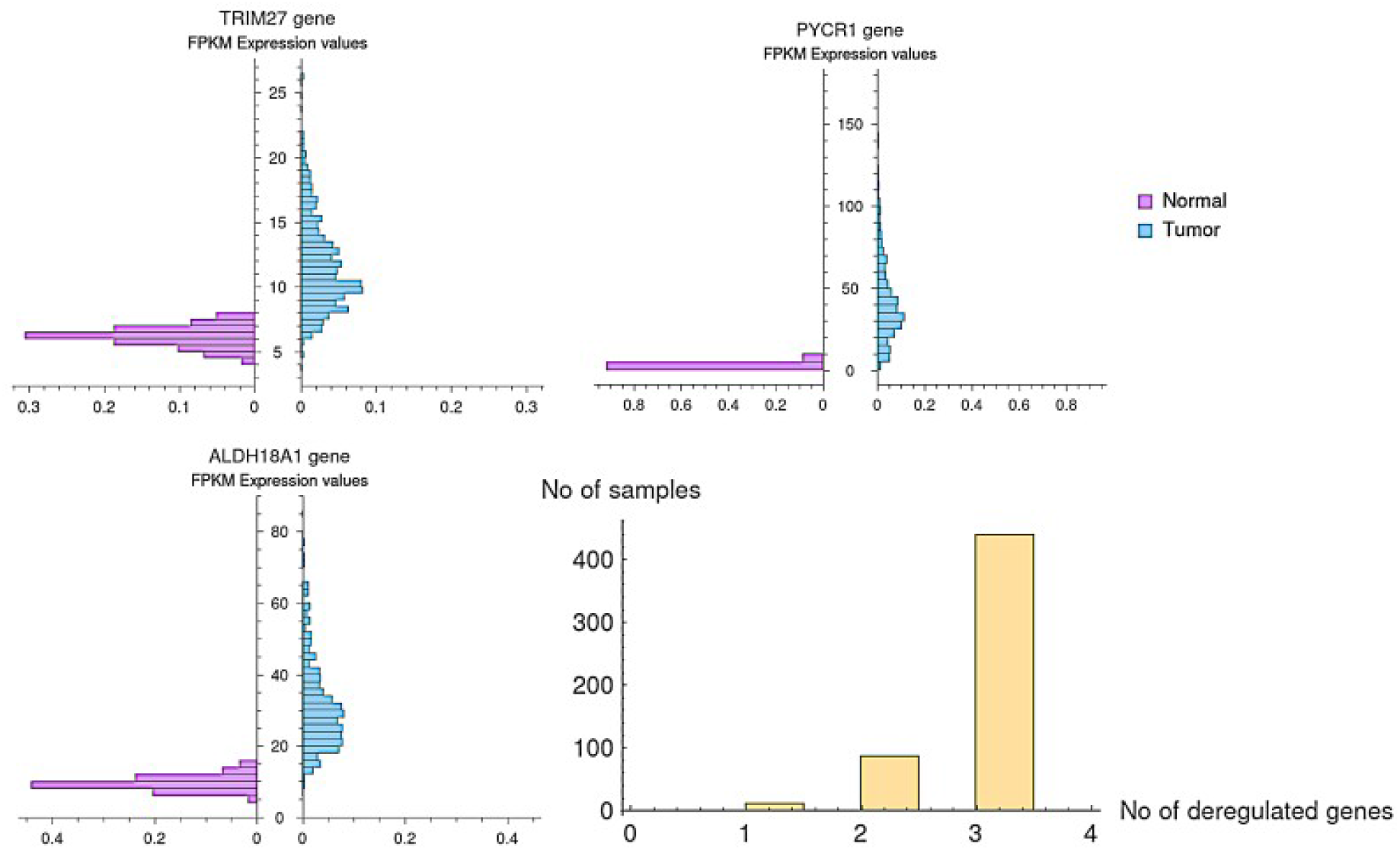
TCGA-LUAD gene expression data for the 3 genes in the panel. There are “normal” intervals for each gene, i.e. by definition, the expression in a normal sample is normal (0). The tumor expression values, on the other hand, could be normal (0) or deregulated (1), that is values above the normal interval. Notice that the last histogram shows that there is at least one deregulated gene for each tumor sample, thus the panel correctly classifies all of the normal samples with {0,0,0}, and all of the tumors, which show at least one deregulated gene.

In the examined class, these 3 genes conform a perfect panel for LUAD. Indeed, notice that the expression values {0,0,0} characterize normal samples, whereas for any tumor sample there is at least one deregulated gene, as shown in the last histogram in **Fig. 2**.

The possibility to avoid false tumor samples (i.e. with expressions {0,0,0}) is related to the existence of genes with very high deregulation frequencies in the tumor population. A single gene panel could be obtained if the gene is deregulated in the whole tumor set. This is what actually happens, for example, with the SCARA5 gene (only normal above class) in colon adenocarcinoma (COAD) **[2]**.

Once examined this particular example, let us come back to the methodology. First, pick up a class of genes (normal concept – only tumor above in the example; at least other nine classes are also possible **[2]**), then construct the binarized table for all genes in the class, then organize genes according to the frequencies in the tumor subset, in descending order. The procedure starts with the highest frequency genes and tries to conform a panel in which the whole tumor population is covered, that is at least one gene in the panel is deregulated for any tumor sample. To that aim, we can use genes that are more or less redundantly deregulated in the tumor set. The PYCR1, ALDH18A1 and TRIM27 trio in the examined example is one of such gene panels, in which we also maximize concurrent deregulations.

We show in Table I the main characteristics of the 3 genes in the panel, according to the Genecards webpage **[13]**. Two of them are related to proline catabolism, a mechanism known to play a crucial role in LUAD **[14]**.

**Table I.** Characteristics of the 3 genes in the perfect panel found for LUAD.

| Gene | Gene characteristics and functions (Genecards) | Deregulation frequency in the tumor subset |
| --- | --- | --- |
| ALDH18A1 | Encodes a mitochondrial enzyme. Plays a crucial role in the de novo biosynthesis of proline | 95 % |
| PYCR1 | Housekeeping enzyme that catalyzes the last step in proline biosynthesis. The protein forms a homopolymer and localizes to the mitochondrion. | 95 % |
| TRIM27 | Plays a role in different processes including proliferation, innate immunity, apoptosis, immune response or autophagy. | 90 % |

In quality of instructive exercise, we may check the performance of the found panel in a different data set. Let us consider the very complete study of a Chinese cohort in **Ref. [15]**. RNAseq data for 51 tumor and 49 control samples for LUAD are provided. We checked that normal intervals for the expressions of the trio of genes may be defined, and that the genes are in the tumor above class (**Fig. 3**). The last histogram in **Fig. 3** shows that the panel provides an exact classification of samples. In fact, the TRIM27 gene is redundant in the present case.

**Fig. 3.**
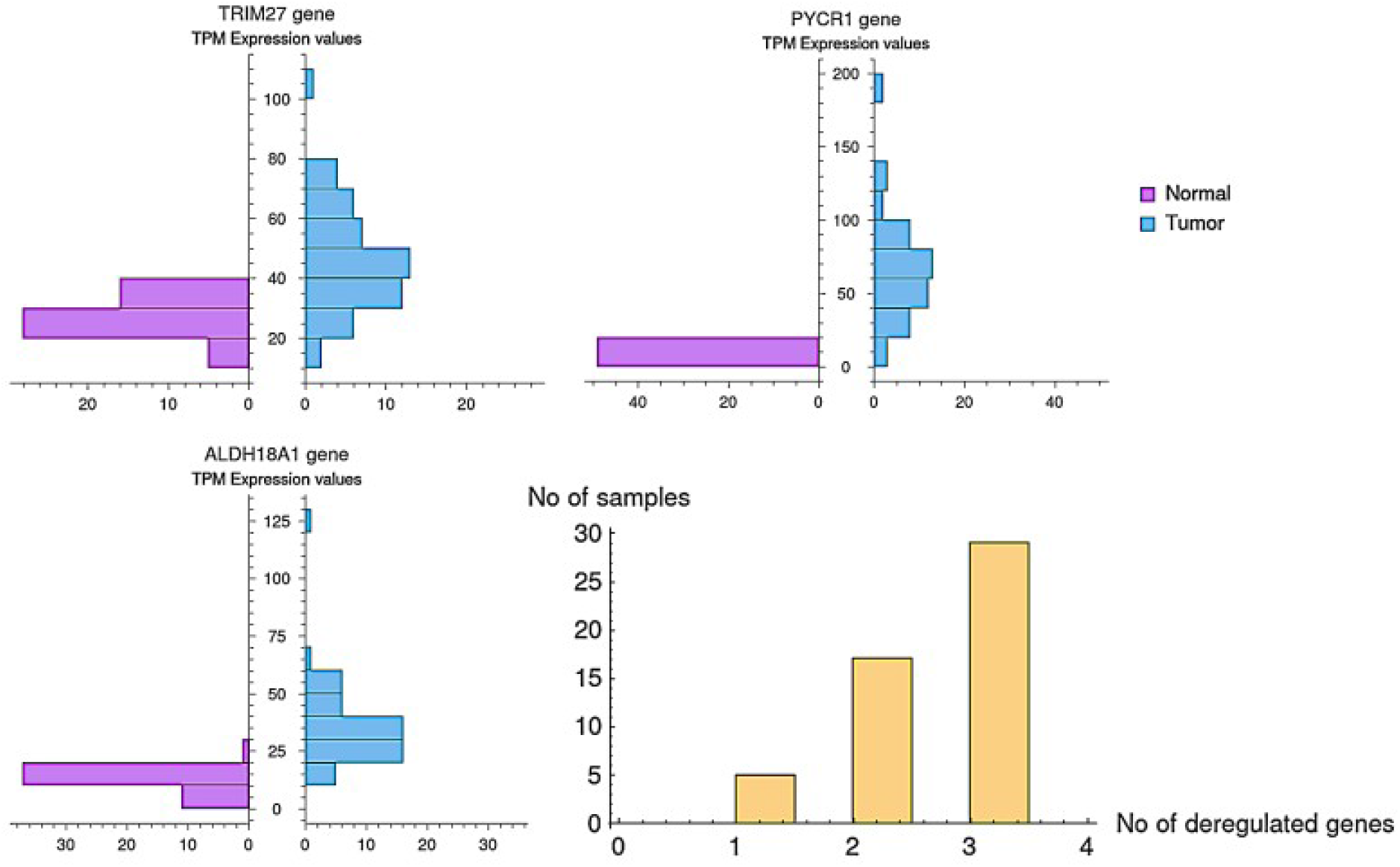
Validation of the 3-gene panel for LUAD by using data of **Ref. [15]** for a Chinese cohort. The expression data shows that the genes remain in the normal concept – tumor above class, whereas the last histogram proves that there is at least one deregulated gene for any tumor sample.

The new data set is interesting not only because the cohort is very different to that one studied in the TCGA portal, but also because the number of samples is one order below. Indeed, the examined TCGA LUAD data contains 59 normal and 535 tumor samples. We may rise the question about the dependence of the number of genes in the panel on the number of tumor samples. The result are summarized in **Fig. 4**. A single gene covers 98 % of the samples in the small set, but only 95% in the larger set. The second gene makes the panel perfect for the small set (the TRIM27 gene is not needed). But in the larger set, there is still a 1% of samples not covered by the 2-gene panel, and we should add the TRIM27 gene. The conclusion is that as the set is enlarged rare variants could emerge. These rare variants are not seen in the smaller sets because their frequency is very low. A point with 99.7% of samples covered by the 3-gene panel in an hypothetical set with 5,000 samples is included in the figure. In this hypothetical scenario, a 4th gene would be needed. The figure also shows that saturation is very rapidly reached. In other words, our statement that a few genes are enough to describe the global state of the GRN seems to be very reasonable and robust (see also **Ref. [2]** for a cross-validation analysis).

**Fig. 4.**
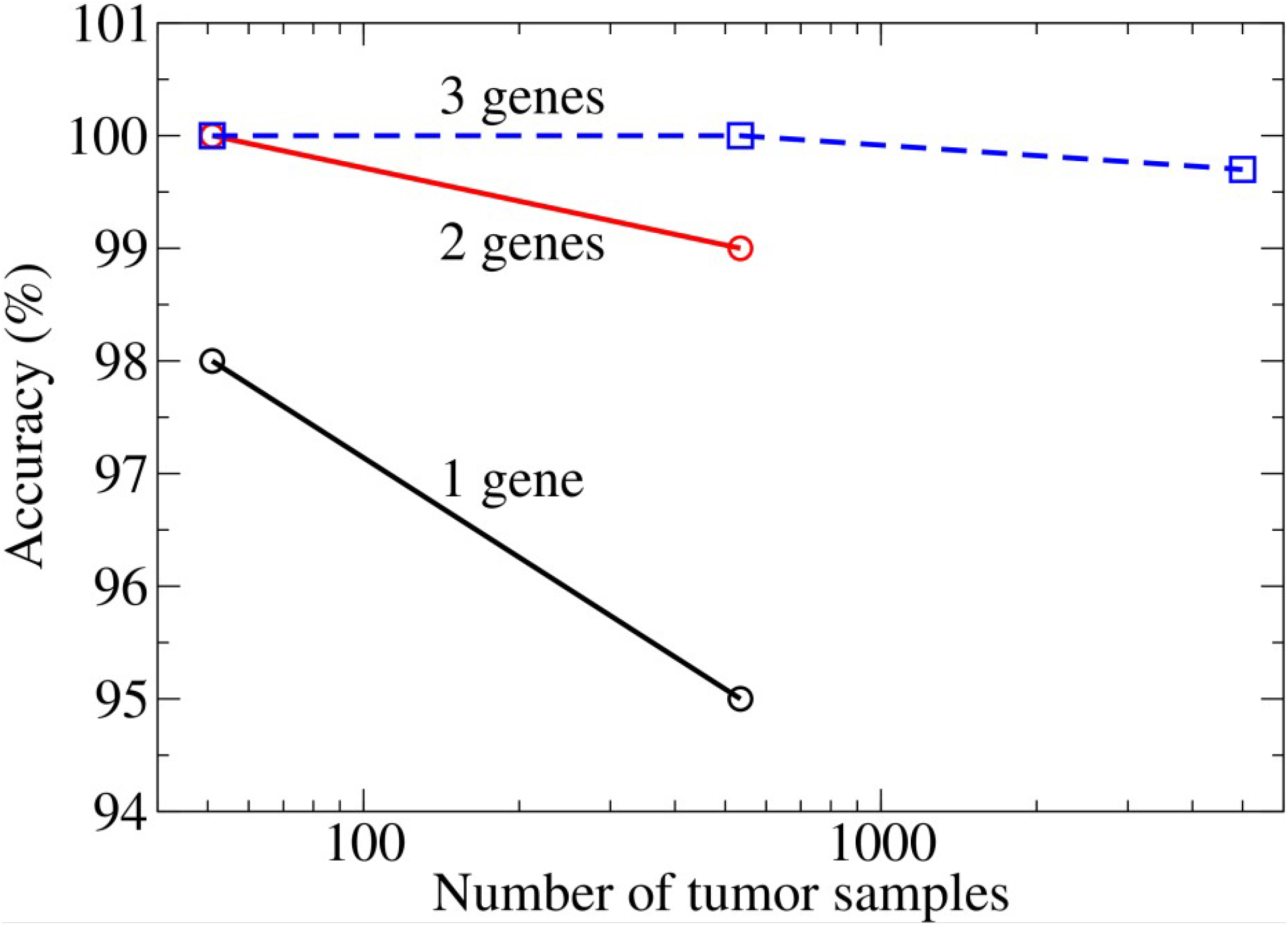
Performance of the one-, two- and 3-gene panels for LUAD as a function of the number of tumor samples. The data from **Ref. [15]** (51 samples), and the TCGA-LUAD data (535 samples) are used. A hypothetical point corresponding to a data with 5,000 samples is also included.

In principle, different panels may be constructed **[2]**. It means that there could be different set of labels for a tumor in a tissue, each of them emphasizing in one or other aspect of the carcinogenic cascade leading to the tumor **[16]**.

We sketch in **Fig. 5** the panels found in **Ref. [2]** for 12 tissues. The *x* axis, as in **Fig. 1**, is the distance between the normal and tumor clouds, whereas the *y* axis shows the lengths of the different panels found for each tissue. The shorter the inter-cloud distance the larger the overlapping between the normal and tumor clouds **[17]**, and thus it becomes harder to classify samples. For close clouds, we need more genes in the panels. And, indeed, the minimal panel in PRAD contains 7 genes, whereas a single gene is enough to perform a perfect classification in COAD, KIRP and UCEC.

**Fig. 5.**
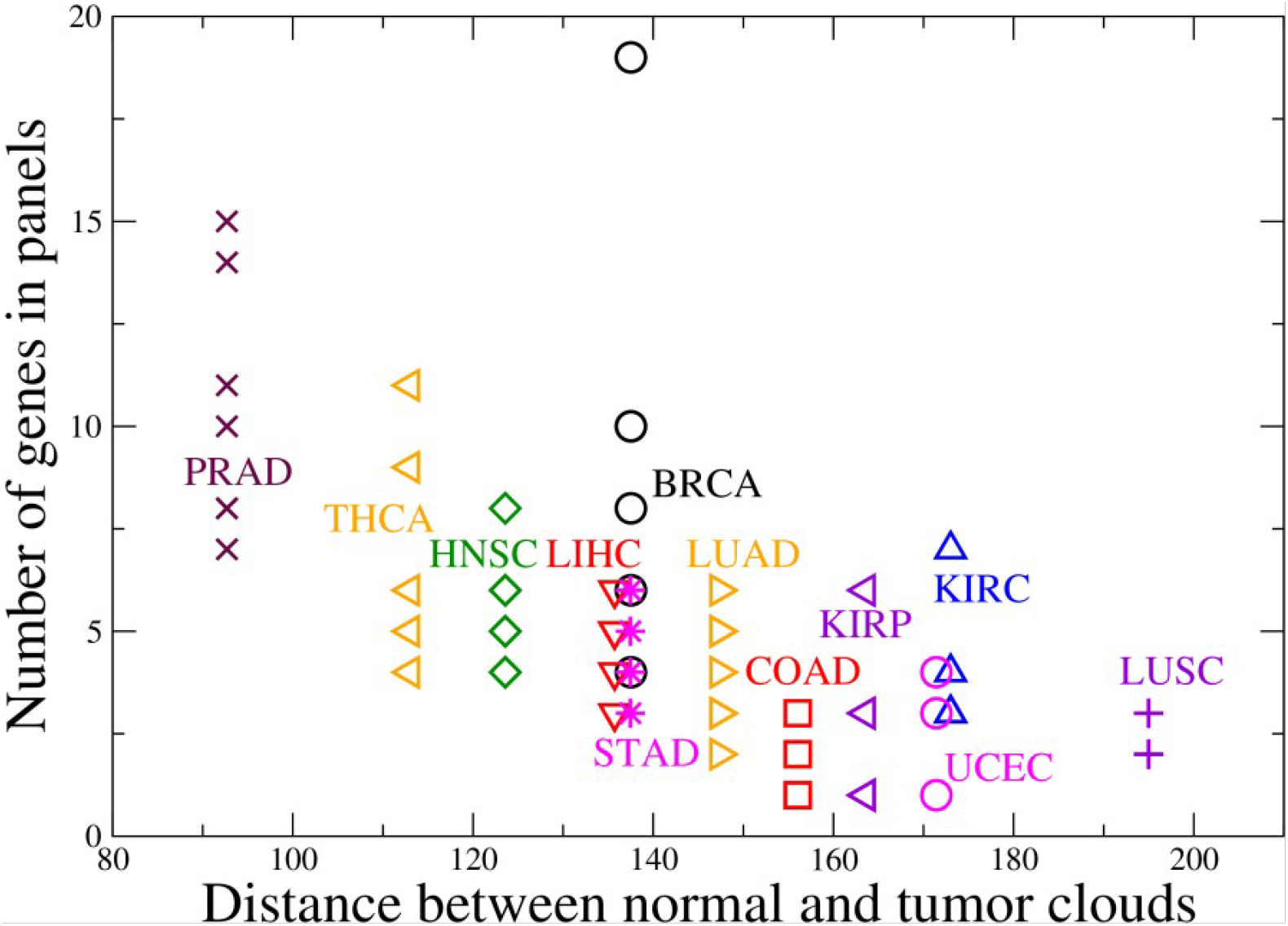
A sketch of the perfect panels found for 12 tissues in **Ref. [2]**. Notice that the minimal panel in PRAD contains 7 genes, whereas 1 gene is enough for a perfect classification in COAD, for example. As expected, tissues with distant less-overlapping normal and tumor clouds are more easily classified.

From an abstract point of view, according to the variance in the dispersion of points, the effective dimensionality of the normal tissue – tumor manifold in gene expression space is around 20 **[17]**. The results of the present paper could be reformulated as the existence of specific directions or sub-manifolds of much lower dimensions, between 1 and 7, the subspaces span by the perfect panels, which allow a complete classification of samples, that is allow to exactly determine the global state of the GRN. We show the inverse correlation between the sub-manifold dimensionality (the length of the minimal panel) and the inter-cloud distance, which is itself a measure of cloud overlapping and dissimilarity between the gene expression profiles in the normal tissue and in the tumor. This last magnitude, the inter-cloud distance in gene expression space, is in fact a global tumor classifier, not often taken into account when studying tumors.

---

**Supplementary Table I.**
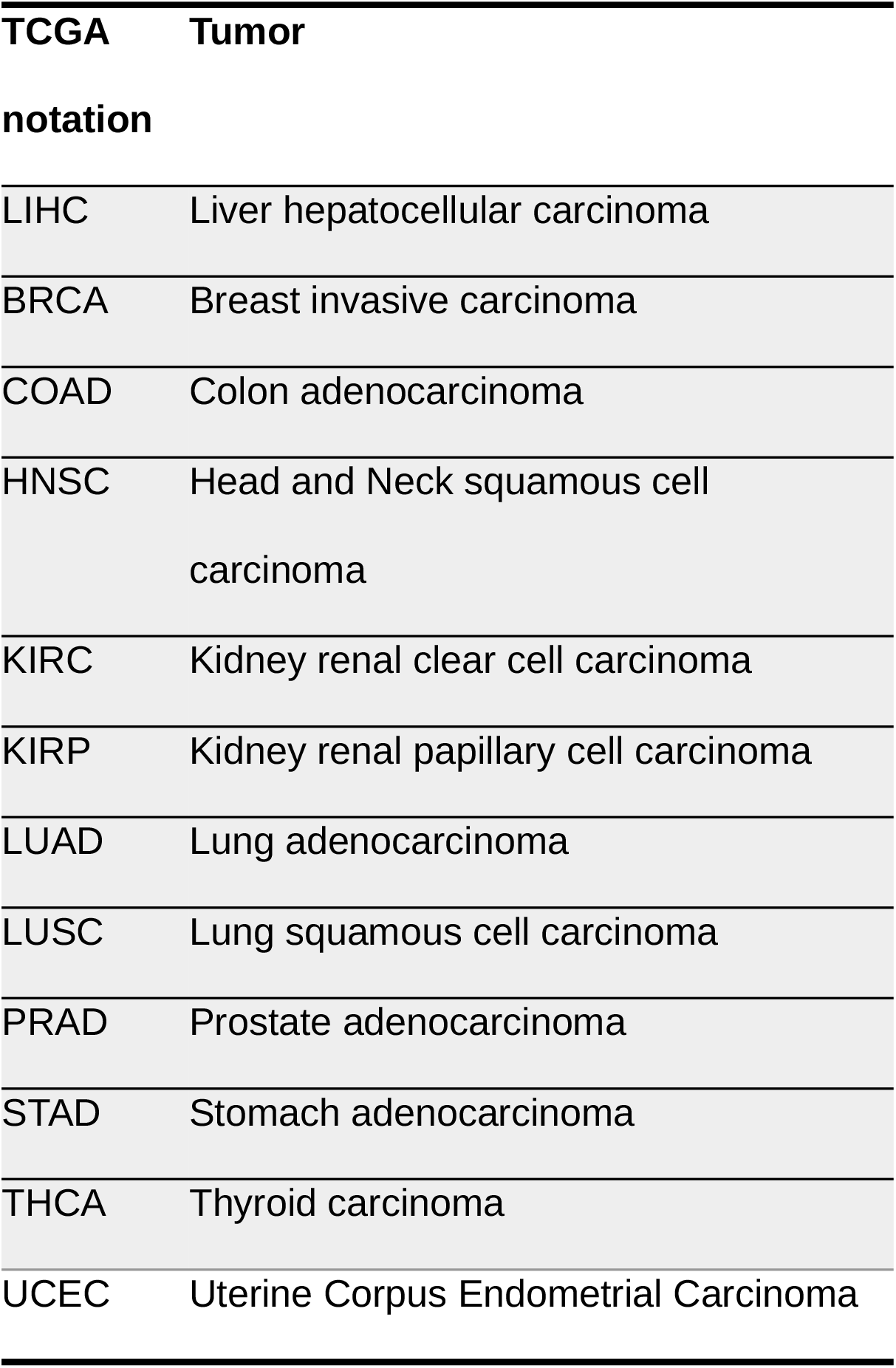
The meaning of the abbreviations used in the TCGA database.

## Acknowledgments

Authors acknowledge the Office of External Activities of the Abdus Salam Centre for Theoretical Physics and the Agency of Nuclear Science and Advanced Technology, Cuba for support.

## Author’s contributions

G.G. ideated the algorithm for the construction of perfect panels and elaborated the inventory for 12 tissues. A.G. wrote an initial draft. Both authors analyzed and interpreted the results, contributed to the manuscript and approved the final version.

## References

[1] Karlebach, G., Shamir, R. Modelling and analysis of gene regulatory networks. Nat Rev Mol Cell Biol 9, 770–780 (2008). 10.1038/nrm2503

[2] Gabriel Gil and Augusto Gonzalez. Perfect genetic biomarkers for cancer from a fresh view of gene dysregulation. biorXiv preprint. 10.1101/2022.07.25.501449 (2022).

[3] Thangavel, K. and Pethalakshmi A. Dimensionality reduction based on rough set theory: A review. Applied Soft Computing 9 (1), 1–12 (2009). 10.1016/j.asoc.2008.05.006

[4] Zhang, S., Xiao, X., Yi, Y. et al. Tumor initiation and early tumorigenesis: molecular mechanisms and interventional targets. Sig Transduct Target Ther 9, 149 (2024). 10.1038/s41392-024-01848-7

[5] Katherine A. Hoadley, Christina Yau, Toshinori Hinoue et al. Cell-of-Origin Patterns Dominate the Molecular Classification of 10,000 Tumors from 33 Types of Cancer. Cell 173 (2), P291-304.e6, 2018. 10.1016/j.cell.2018.03.022

[6] Malone, E.R., Oliva, M., Sabatini, P.J.B. et al. Molecular profiling for precision cancer therapies. Genome Med 12, 8 (2020). 10.1186/s13073-019-0703-1

[7] Durães, C., Pereira Gomes, C., and Costa, JL., and Quagliata, L. (2022), Demystifying the Discussion of Sequencing Panel Size in Oncology Genetic Testing. European Medical Journal 7 (2), 68–77. 10.33590/emj/22C9259

[8] The Cancer Genome Atlas Research Network. Comprehensive molecular profiling of lung adenocarcinoma. Nature 511, 543 – 550 (2014). 10.1038/nature13385

[9] Gonzalez, A., Leon, D. A., Perera, Y. & Perez, R. On the gene expression landscape of cancer. PLOS ONE 18, e0277786 (2023). 10.1371/journal.pone.0277786

[10] Duhan, N., Sharma, A. K. & Bhatia, K. K. Page Ranking Algorithms: A Survey. 2009 IEEE International Advance Computing Conference, 1530–1537 (2009). 10.1109/iadcc.2009.4809246

[11] Gonzalez, A., Nieves, J., Leon, D. A., Bringas Vega, M. L. & Sosa, P. V. Gene expression rearrangements denoting changes in the biological state. Scientific Reports 11, 8470 (2021). 10.1038/s41598-021-87764-0

[12] Nieves J, Gonzalez A. The Geometry of Normal Tissue and Cancer Gene Expression Manifolds. Acta Biotheor. 2024 Jul 9;72(3):9. 10.1007/s10441-024-09483-z

[13] The GeneCards Suite: From Gene Data Mining to Disease Genome Sequence Analyses. Stelzer G, Rosen R, Plaschkes I, et al. Current Protocols in Bioinformatics (2016), 54:1.30.1 – 1.30.33. 10.1002/cpbi.5

[14] Liu, Y., Mao, C., Wang, M. et al. Cancer progression is mediated by proline catabolism in non-small cell lung cancer. Oncogene 39, 2358–2376 (2020). 10.1038/s41388-019-1151-5

[15] Jun-Yu Xu, Chunchao Zhang, Xiang Wang, et al. Integrative Proteomic Characterization of Human Lung Adenocarcinoma. Cell 182, 245–261 (2020). 10.1016/j.cell.2020.05.043

[16] Roberto Herrero, Jean Pierre Gomez, Gabriel Gil and Augusto Gonzalez. Deregulation cascades in carcinogenesis. BiorXiv preprint 2023.12.28.573550 (2023). 10.1101/2023.12.28.573550

[17] Augusto Gonzalez, Frank Quintela, Dario A. Leon, Maria Luisa Bringas-Vega, and Pedro A. Valdes-Sosa. Estimating the number of available states for normal and tumor tissues in gene expression space. Biophysical Reports 2, 100053 (2022). 10.1016/j.bpr.2022.100053

